# Molecular mechanism of nitrogenase sequestration by a P_II_-protein couple

**DOI:** 10.64898/2025.12.07.692841

**Authors:** Nevena Maslać, Pauline Bolte, Marie-Caroline Müller, Tristan Wagner

## Abstract

The microbial process of N_2_-fixation is crucial for the planetary nitrogen cycle and biosphere, but requires substantial cellular energy resources. Here, we solved how energy-limited anaerobes regulate on-demand N_2_-fixation by sequestering their nitrogenase through the P_II_-nitrogen regulatory proteins NifI_1_ and NifI_2_. The nitrogenase was directly isolated from a methanogenic archaeon, with NifI proteins tightly bound. The crystal structure of the inhibited form, refined to 2.3-Å resolution, reveals a supercomplex in which three NifI_1,2_ units made of a NifI_1,2_ heterohexamer captured three nitrogenases via tentacular T-loops. Additional structural information confirmed that NifI_1,2_ sits at the nitrogenase reductase-binding site, preventing N_2_-reduction. The presence of MgATP and 2-oxoglutarate releases NifI_1,2_ from the nitrogenase core via a conformational switch of the T-loops, provoking a steric repulsion and loss of contacts. The overall molecular depiction corroborates previous genetic, biochemical, and biophysical experiments, proposing that NifI_1,2_ disrupts the dynamic nitrogenase-reductase association, thereby interfering with electron delivery for N_2_-fixation and preventing ATP consumption. While ligand-binding mode and the T-loop conformational switch are expected to be conserved among NifI_1,2_-utilisers, the association mode with the nitrogenase comes in different flavours as a few substitutions in NifI_2_ break NifI_1,2_ intramolecular dimerisation in *Methanosarcinales* species, readjusting the supercomplex without altering the inhibition mechanism. With the NifI_1,2_ allosteric control dependent on the alarmone 2-oxoglutarate, anaerobes can effectively balance nitrogen-acquisition versus energy-expenditure, a regulatory switch that might be primitive and has been progressively lost in non-energy-limited aerobes, but could inspire biotechnological engineering to optimise ammonia bioproduction.

## Introduction

Biological N_2_-fixation catalysed by nitrogenases feeds the biosphere with bioavailable nitrogen in the form of ammonia. The nitrogenase splits the N_2_ triple bond with the FeMo-cofactor (FeMoco), a metallocluster that catalyses the most energetically expensive reaction known in biology (Supplementary Fig. 1a). The baseline energy demand of the most efficient nitrogenase system has been long established as 16 ATP consumed per N_2_ reduced in an ideal case scenario (1, 2). However, recent revisions have increased this requirement to approximately 25 ATP consumed per N₂ (3), a value that still excludes the considerable redox investment required to supply low-potential electrons to the enzyme complex. The chemical energy is required for the electron delivery from the nitrogenase reductase (NifH) to the nitrogenase core (organised in a tetramer of 2(NifDK)). It results in the hydrolysis of two ATP per injected electron and the undesirable spontaneous proton reduction by the FeMoco under catalytic turnover (Supplementary Fig. 1a). While this energetic toll can be effectively offset in non-energy-limited organisms, such as symbiotic species or aerobically respiring bacteria (as the model organism in the study of nitrogenase, *Azotobacter vinelandii*), it may pose a substantially greater burden for anaerobes, whose energy supply is constrained by the thermodynamic limits of their catabolic reactions and the limited substrate availability. For these microbes, tight regulation of the nitrogenase is necessary to prevent undesired ATP hydrolysis, particularly in diazotrophic methanogenic archaea that thrive at the thermodynamic limits of Life (4–6). Methanogens are strict anaerobes that play a key role in two biochemical cycles: carbon, contributing to half of yearly atmospheric methane emissions, and nitrogen, by actively fixing N_2_, and even being the prominent diazotrophs in some ecological niches (7–10). The laboratory of John A. Leigh provided decades of seminal work on how methanogens regulate their N_2_-fixing system by using the genetically tractable model *Methanococcus maripaludis* (11–13) and discovered that the nitrogenase is regulated at the transcriptional level by the repressor NrpR (14), and at the post-transcriptional level through direct control by the P_II_-nitrogen regulatory proteins NifI_1_ and NifI_2_ (15–17). P_II_-proteins are key regulators of the nitrogen metabolism and interfere with cellular nitrogen trafficking, such as ammonium transportation (18, 19), ammonium fixation by glutamine synthetase (20–23) or through the ornithine-ammonia cycle (24). They all form homotrimers with protruding loops (named B-, C-, and T-loops) that serve as the binding sites for MgATP and 2-oxoglutarate (2OG) at their base. Once MgATP and 2OG are bound to the interdimeric clefts, a conformational switch displaces the T-loop, modifying the functional properties of the P_II_-protein (Supplementary Fig. 1a (25–29)). Therefore, the P_II_-protein is directly under the control of cellular energy and nitrogen levels (25–29). In *M. maripaludis*, the inhibitory P_II_-proteins NifI_1,2_ bind to the nitrogenase, forming a large assembly (500-700 kDa). The assembly disaggregates upon the addition of ATP and 2OG, restoring nitrogenase activity by switching the regulator to an off-state (16). Further experiments showed that NifI_1,2_ inhibition cannot be achieved with either NifI_1_ or NifI_2_ alone and that variants lacking the T-loop are less effective at interacting with the nitrogenase. Moreover, competitive experiments using the reductase NifH led to a proposed scenario in which NifI_1,2_ would inhibit N_2_-fixation by interfering with NifH (30).

In this work, we relied on the hyperthermophile *Methanocaldococcus infernus* to decipher the molecular basis behind NifI_1,2_ inhibition of nitrogenase, using structural snapshots of the different actors involved. While it also belongs to the marine hydrogenotrophic *Methanococcales* order, as *M. maripaludis*, *M. infernus* is a hyperthermophile capable of fixing N_2_ up to 91.3 °C, harbouring a remarkable hyperthermostable nitrogenase (31). We previously showed that the FeMoco-containing nitrogenase naturally forms a stable complex with endogenous NifI_1,2_ when isolated from the lysate (31). The complex exhibited behaviour similar to that of *M. maripaludis*, with a 10% activity for the as-isolated inhibited form, which restored full N_2_-fixation rates upon addition of 10 mM 2OG (MgATP being required in the activity assay). The measured K_0.5_ of 1.4 mM for 2OG in the *M. infernus* system is also similar to the K_0.5_ of 6 mM measured in *M. maripaludis*. To unveil how NifI_1,2_ binds to NifDK, the isolated complex was anaerobically purified (Supplementary Fig. 1b) and crystallised, and the structure was solved by X-ray crystallography.

### As-isolated NifDKI_1,2_ forms a supercomplex

The NifDKI_1,2_ structure, refined to 2.3 Å resolution (Supplementary Table 1), adopts an 863 kDa supercomplex architecture, in agreement with the assembly reported in *M. maripaludis* (16). The overall assembly can be decomposed as follows: a trimeric arrangement of the nitrogenase lies in the centre, flanked by three NifI_1,2_ units (Fig. 1a, Supplementary Fig. 2a). The NifI_1,2_ unit is constituted by four NifI_1_ and two NifI_2_ that dimerise through NifI_2,_ forming a 2(2NifI_1_NifI_2_) oligomer. In this form, the NifI_1,2_ unit can capture two nitrogenases on each side, similar to the way immunoglobulin G binds to its target. Since the nitrogenase itself is a NifDK homodimer, this arrangement could, in principle, support the formation of indefinite filamentous assemblies. However, in contrast to the Shethna protein II, which elongates in filaments (32, 33), the kinked shape of NifI_1,2_ closes the oligomerisation into a triangular architecture (Fig. 1a, Supplementary Fig. 2a).

**Fig. 1.**
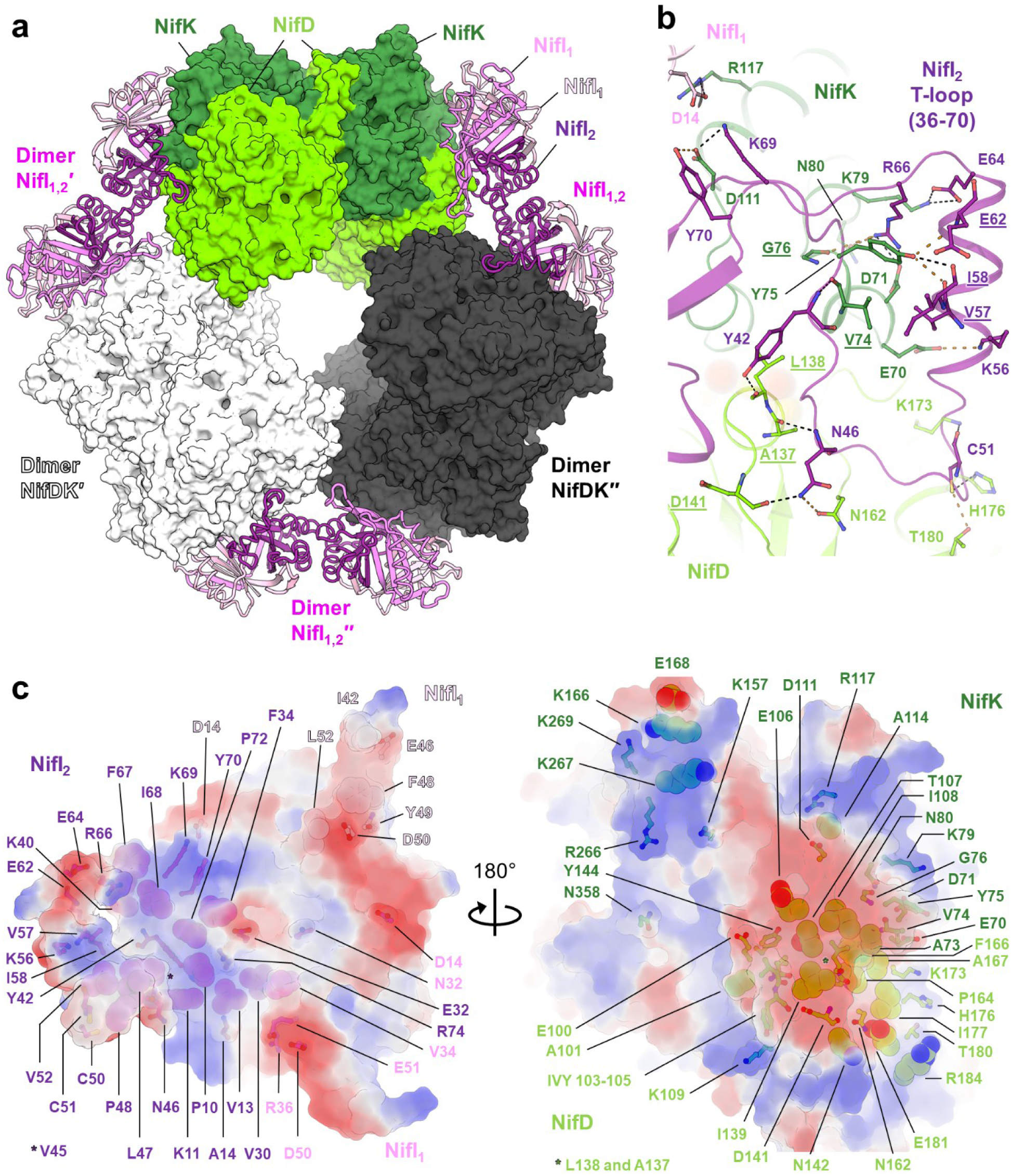
*M. infernus* NifDKI_1,2_ supercomplex. **a,** Overall view of the NifDKI_1,2_ supercomplex, in which NifDK is displayed as surfaces and NifI_1,2_ as cartoons. **b**, Close-up of the interaction of NifI_2_ T-loop with NifDK. Proteins are shown in cartoons, with residues interacting via hydrogen bonds (displayed by dashed lines, with orange ones being at a distance of 3.5-4.0 Å) highlighted in sticks. Residues interacting only via the main chain are underlined. **c**, Electrostatic surface complementation between NifI_1,2_ (left) and NifDK (right), ranging from negatively (red) to positively (blue) charged. Residues involved in hydrogen bonds and Van der Waals interactions are shown in sticks and balls, respectively.

Most of the interaction between the NifI_1,2_ units and the nitrogenase occurs at the NifI_2_ T-loop (residues 36-70), which wraps around NifD and NifK (Fig. 1b, Supplementary Figs. 3 and 4). Additional contacts are mediated by the NifI_1,2_ trimeric core, whereas the NifI_1_ T-loops (residues 36-51, Supplementary Figs. 3) contribute only a few hydrogen bonds (Supplementary Fig. 2b) and Van der Waals contacts (Fig. 1c). Assembly stabilisation is mediated by an extensive interface driven by electrostatic complementarity and hydrophobic interactions at its core (Fig. 1c). The nitrogenase stacked in this trimeric arrangement engages salt-bridge contacts through NifD, ensuring a locked stable spatial configuration while avoiding clashes (Supplementary Fig. 2c, Supplementary Fig. 4). The overall structure of nitrogenase remains unchanged upon NifDK stacking, showing no deviations from the stand-alone NifDK (Supplementary Fig. 5a). The different metallocofactors (the magnesium atom at the NifK-NifK′ dimer interface, the electron transferring P-cluster, and the catalyst FeMoco) are present at full occupancy (Supplementary Fig. 5b). The FeMoco is found in a classic resting state, well-described in this type of enzyme, including in *M. infernus* (Supplementary Fig. 5b)(31, 34, 35). The P-cluster has been modelled as a fully reduced state and might contain partially occupied oxidised forms (i.e., P^1+^ and P^2+^), which cannot be attributed with confidence at such resolution (Supplementary Fig. 5b). The presence of the mild reductant dithiothreitol over the purification and crystallisation could have safeguarded the nitrogenase from oxidative damage. The preserved structural integrity of the metallocofactors corroborates the observation of high activity recovery upon NifI_1,2_ release (31). To understand the mechanism of NifI_1,2_ release, we further isolated NifI_1,2_ from the nitrogenase by adding MgATP and 2OG (see methods, Supplementary Fig. 1b), purified it to homogeneity and co-crystallised it with its ligands.

### Ligand-induced switch-off state of NifI_1,2_ disrupts association with NifDK

The crystal structure of NifI_1,2_ in a holo state (i.e., in complex with MgATP and 2OG), refined to 2.7-Å resolution (Supplementary Table 1), reveals the same NifI_1,2_ unit as previously described in the NifDKI_1,2_ supercomplex (Fig. 2a, Supplementary Fig. 6). A detailed analysis shows that within each heterotrimers, NifI subunit binds tightly to the two others through extensive hydrophobic contacts locked by salt bridges, reinforcing the assembly (Supplementary Fig. 7a). The NifI_2_ dimerisation interface is constituted by helices α1 and α3 that form a four-helix bundle. Despite the restrained contact area, the dimeric interface is highly stabilised through hydrophobic contacts surrounded by salt bridges (Supplementary Figs. 3 and 6). Both NifI_1_ and NifI_2_ subunits harbour a MgATP and 2OG locked in the interdimeric cleft formed by the three protrusions B, T, and C-loops, as previously seen in P_II_-protein homologues (Fig. 2a-b, Supplementary Figs. 6 and 7b). Ligand interaction with the protein matrix and the residues involved in binding are almost perfectly conserved between the three different sites formed by the hybrid combination of Nif_2_/NifI_1_, NifI_1_/NifI_2_ and Nif_1_/NifI_1_ (Fig. 2b, Supplementary Fig. 3). When compared to the close homologue GlnK_2_ from *Methanothermococcus thermolithotrophicus* only a few positions directly interacting with ATP or 2OG are substituted or slightly shifted (Fig. 2b). This subtle readjustment of bonds might tune ligand affinity, explaining the relatively high K_0.5_ for 2OG (mM range) in NifI_1,2_ compared to the high-affinity GlnK (µM range)(36). The geometry of the ATP and 2OG, both coordinated by the Mg^2+^, is identical to that observed in the GlnK protein (Fig. 2b, Supplementary Figs. 1a and 7b). Such conservation suggests that the conformational switch provoked by ligands, common to all P_II_ proteins, would act similarly in NifI_1,2_, thereby displacing the T-loops and dramatically altering their functional properties. Indeed, a comparison of the apo and the holo states shows a major displacement in the T-loops of NifI_1_ and NifI_2_, while B- and C-loops remain mostly static (Fig. 2a, Supplementary Figs. 6 and 7b). This displacement switches the T-loops from a flat conformation in the apo state to an extended conformation protruding from the NifI_1,2_ core at a ≈60° angle (ranging from 45 to 75°). In the NifI_2_ holo form, the long T-loop is mostly disordered, but still shows drastic movement at its base. When the holo state is superposed on the apo state of the NifDKI_1,2_ complex, NifI_1_ T-loops repositioning generates substantial steric clashes with NifDK, while NifI_2_ T-loops base shows an expected disengagement that would cause a break of the hydrogen bond network and Van der Waals interactions (Fig. 2c).

**Fig. 2.**
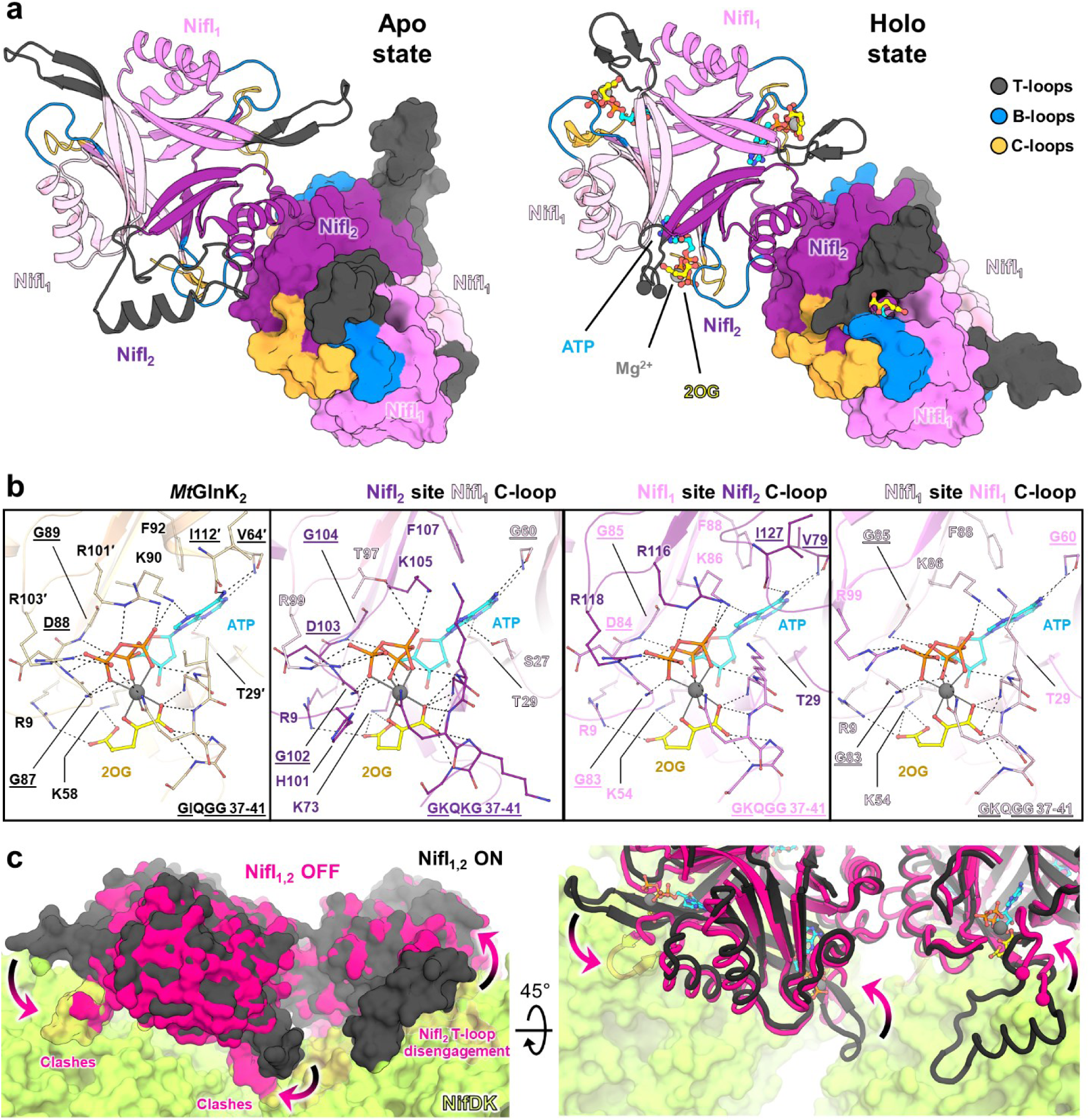
NifI_1,2_ conformational changes upon ligand binding. **a**, NifI_1,2_ apo and holo overall view in which one NifI_1,2_ trimer is shown as a cartoon and the other as a surface. In the apo structure, one NifI_1_ (in light pink surface) has a disordered unmodelled T-loop. In the holo structure, both NifI_2_ have a disordered, unmodelled T-loop, as shown by the cut indicated by balls in the cartoon model. Ligands are shown in balls and sticks. T, B, and C-loops involved in ligand binding are differently colored. **b**, Close-up view of the ligand binding site. Residues interacting with the ligands via hydrogen bonds (displayed in dashed lines) are shown as sticks. Residues interacting only via the main chain are underlined. A canonical phenylalanine interacting with the ATP is also shown (i.e., F92 for *Mt*GlnK_2_). Mg^2+^ is presented as a grey ball. GlnK_2_ from *Methanothermococcus thermolithotrophicus* (*Mt*GlnK_2_, PDB 7P50) was added as a comparison (27). **c**. Steric repulsion between NifI_1,2_ and NifDK induced by ligand binding. In the left panel, all proteins are represented as surfaces, while the right panel shows NifI_1,2_ as cartoons with ligands as balls-and-sticks, coloured as in panel b. Arrows depict T-loop movements.

To summarise, the overall conformational changes induced by MgATP and 2OG binding readjust the NifI_1,2_ T-loops to an off-state incapable of binding NifDK due to steric repulsion and the loss of major contacts. To resolve how NifI_1,2_ inhibits N_2_-reduction, we investigated the binding of the nitrogenase to the reductase unit during electron delivery.

### NifI_1,2_ prevents NifH association by occupying its binding site

The pool of isolated NifDK (31) was combined with purified NifH (37) and co-crystallised with MgADP and AlF_4_^-^ to stabilise the complex in a conformation that precludes ATP hydrolysis, mimicking a transitional state immediately before electron transfer (Supplementary Fig. 1)(38–40). The crystal structure of the NifDKH complex refined to 2.6-Å resolution (Supplementary Table 1) presents the nitrogenase capped by two NifH dimers (simplified here as NifH) containing the MgADP-AlF_4_^-^ (Fig. 3a, Supplementary Figs. 8-9). It is worth noticing that one of the NifH dimers exhibits a high B-factor (Supplementary Fig. 8a), a wobbly position not maintained by the crystalline packing, which supports the proposal that only one NifH should be bound at a time during turnover conditions (41). The NifH position on the nitrogenase fits the one described for the homologue from *A. vinelandii* (38), which was also trapped with a nucleotide analogue mimicking hydrolysed ATP. In this position, the NifH dimer aligns with the pseudo C2-symmetry axis of NifDK and places electron-transferring metalloclusters front to front (Fig. 3b, Supplementary Fig. 8b). The [4Fe-4S] cluster coordinated by NifH and the P-cluster are separated by a distance of 14.5-Å allowing efficient electron transfer as in NifDK from *A. vinelandii* (Fig. 3a, Supplementary Fig. 8c). Only local minor structural differences that do not impact the overall NifDK architecture can be observed on NifDK upon complexation with NifH (Supplementary Fig. 5a). Due to the resolution and an inherent crystallography defect (see Materials and Methods) the redox state of the P-cluster cannot be accurately assigned and was modelled as a mixture of oxidised and reduced states (i.e., P^2+^ and P^0^, respectively), while the FeMoco is in its classic resting state (Supplementary Fig. 5b).

**Fig. 3.**
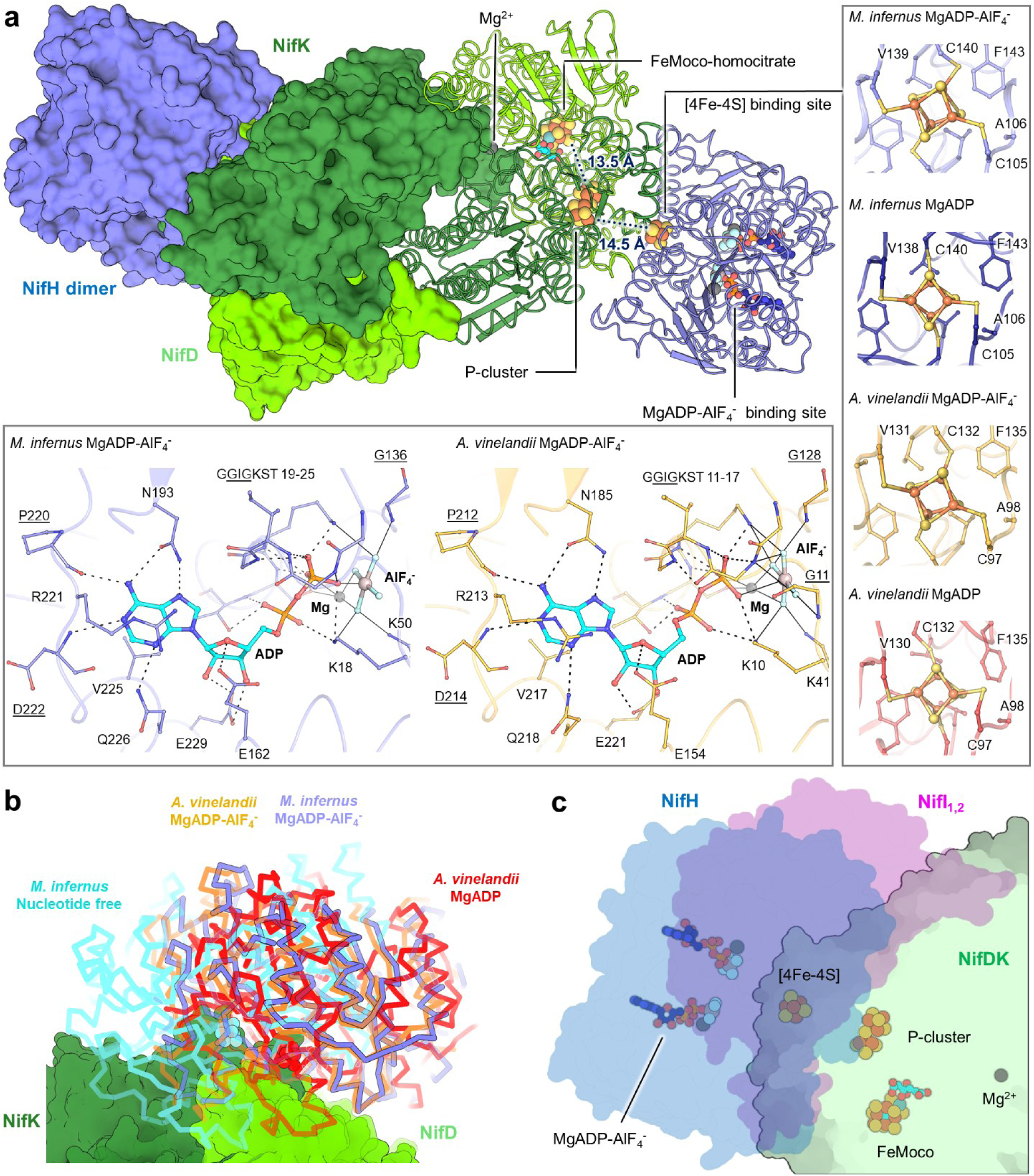
NifDKH complex and NifI_1,2_ inhibition by competition. **a**, NifDKH complex, with half shown as a surface and the other half as a cartoon. Ligands are shown as balls and sticks. Framed inlets are close-ups of the [4Fe-4S] and MgADP-AlF_4_^-^ binding sites. Residues involved in hydrogen bond contacts (dashed lines) and Mg/AlF_4_^-^ interactions (straight lines) are shown as balls and sticks. Underlined residues interact solely via the main chain. **b**, Superposition of NifDKH complexes of *A. vinelandii* on NifDKH from *M. infernus*, in which only NifDK from *M. infernus* is shown as a surface and NifHs as a ribbon. The structure from *M. infernus* presents a NifH position similar to the complex of *A. vinelandii* with MgADP-AlF_4_^-^. Used PDB codes are described in Supplementary Fig. 8. **c**, Superposition of NifDKH and NifDKI_1,2_ on NifDK core, highlighting the colocalisation of NifI_1,2_ and NifH. Ligands are represented and colored as in panel **a**.

A detailed comparison between *M. infernus* system with the well-studied bacterial counterpart led us to pinpoint its highly conserved traits: (*i*) NifH recognizes NifDK mostly through an extensive electrostatic interaction with a positively charged centre flanked by negatively charged discs that adequately fit NifDK surface (Supplementary Fig. 9); (*ii*) Upon the addition of AlF_4_^-^, MgADP-NifH follows a conformational movement due to readjustment of structural determinants leading to a modification of the [4Fe-4S] cluster environment (Fig. 3a, Supplementary Fig. 8d-e); (*iii*) the binding mode of the ligands and their coordination in NifH is also extremely conserved (Fig. 3a). This high structural conservation infers that the same electron delivery mechanism described and detailed for decades (1, 35, 41–44) applies to the archaeal system of *M. infernus*. Therefore, it is expected that the sequential choreography leading to NifH-NifDK recognition, ATP hydrolysis coupled to electron delivery, and dissociation upon phosphate release follow the same course (Supplementary Fig. 1a), with each of these steps being potentially blocked in the NifDKI_1,2_ supercomplex.

A superposition of NifDKH complexes trapped in *A. vinelandii* on the trimeric form of NifDK imposed by NifI_1,2_ confirmed that none of the NifH-bound states clash with the nitrogenase trimeric assembly itself (Supplementary Fig. 8d). However, the superposition of NifDKH and NifDKI_1,2_ from *M. infernus* shows that the dimeric NifH and heterotrimeric NifI_1,2_ share the same binding site (Fig. 3c). The co-localisation would force the competition between the two for binding on the nitrogenase core. However, given the larger contact surface of NifDK-NifI_1,2_ (2,477 Å²) compared to NifDK-NifH (1,894 Å²), and considering that the NifH dimer must interact dynamically with NifDK for efficient turnover, it can be inferred that active NifI_1,2_ will outcompete NifH. Consequently, the ferocious binding and stabilisation of NifI_1,2_ on NifDK prevent NifH from accessing its binding site and transferring electrons to nitrogenase, thereby saving ATP and redox power when cellular nitrogen is sufficient (Fig. 4).

**Fig. 4.**
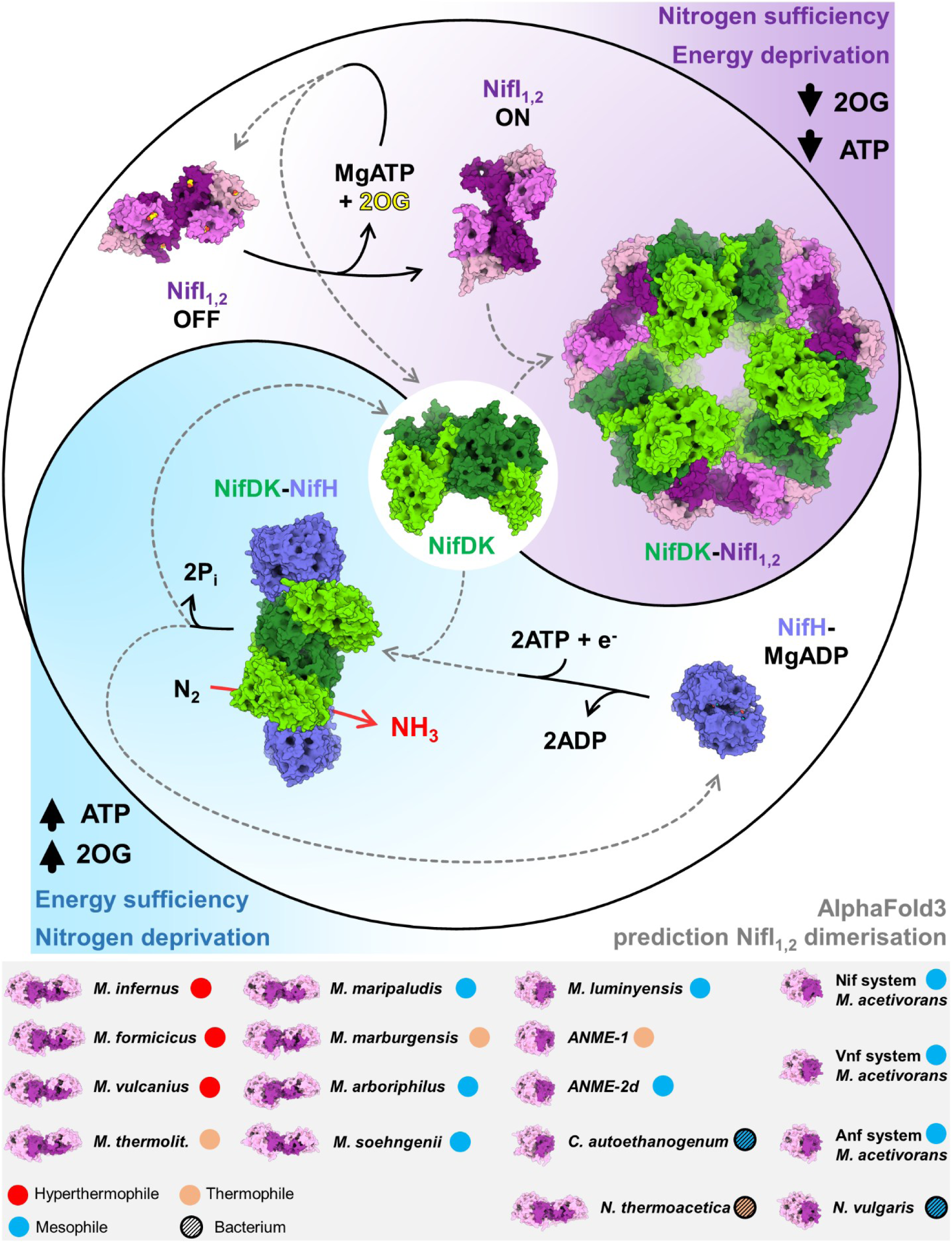
Proposed mechanism governing the regulation of N_2_-fixation by NifI_1,2_. NifH-MgADP complex comes from PDB 8Q5W (37). The bottom panel presents computationally generated NifI_1,2_ models from methanogens, methanotrophs, and anaerobic bacteria (see Supplementary Fig. 4 for complete names).

### Conservation and variances in NifI_1,2_ regulation

Since the residues constituting the MgATP/2OG binding pocket are mostly conserved in methanogens, methanotrophs and bacterial homologues based on sequence alignment (Supplementary Fig. 3), it is expected that NifI_1,2_ homologues would conserve ligand specificity and share the mechanism of the T-loop conformational switch. However, it does not warrant that the ligand binding mode and NifDK supercomplex organisation will be identical in other NifI_1,2_-containing microbes. In fact, a recent report on the NifDK-NifI_1,2_ supercomplex of *Methanosarcina acetivorans* shows a mixture of bound ADP and 2OG at the different NifI_1,2_ sites in the isolated complex (45). In that case, the 2OG is bound independently of a Mg-nucleotide in the ATP binding site. More strikingly, the overall NifDKI_1,2_ complex is differently organised, with a tightly packed trimer of nitrogenase sequestered by a NifI_1,2_ heterotrimer that does not undergo dimerisation. The absence of NifI_2_-NifI_2_ dimerisation leads to readjustments in contacts between NifDK and NifI_1,2_, affecting the overall assembly. However, the 2xNifI_1_-NifI_2_ basic heterotrimer, the trimeric NifDK sequestration and the inhibition mechanisms by interfering with the NifH binding site remain conserved. The dimerisation of NifI_2_ in *M. infernus* is unlikely to be a hyperthermophilic trait, as computational modelling of homologues proposes that a similar organisation is widespread in other hydrogenotrophic methanogens (e.g., *Methanobacteria*) and bacteria (e.g., *Firmicutes*). (Fig. 4).

The difference in the supra-organisation of NifDKI_1,2_ between these two methanogens might confer variation in the inhibitory response, concealing a more intricate regulatory system in *Methanosarcinales*. This concept is further reinforced by the observation that the NifI_1,2_ inhibitory effect is more drastic in *Methanococcales* (i.e., 10% basal nitrogenase activity when fully inhibited by NifI_1,2_ in *M. infernus*/*M. maripaludis* (15, 31)) than in *Methanosarcinales* (i.e., 43% basal activity in *M. acetivorans* when fully inhibited by NifI_1,2_ (45)). The same difference in regulation has already been observed in the glutamine synthetases from *Methanococcales* (46) and *Methanosarcinales* (20, 23, 25, 47). Here, the NH_3_-fixing enzyme in *Methanococcales* is inactive as-isolated and requires only 2OG to restore its full activity, whereas in *Methanosarcina mazei* the same enzyme is additionally regulated by another P_II_-protein (i.e., GlnK_1_) and a small protein (i.e., sP26 (48)) that fine-tunes its activity rate. The simplification of nitrogen assimilation regulation described here could be extended to methanogens with an exclusive hydrogenotrophic lifestyle (Fig. 4), which constrain metabolic versatility and energy yield. It may suggest a primitive system that relies solely on the master regulator 2OG (49) and ATP, in line with the proposal that the nitrogenase of *Methanococcales* is ancient, if not the progenitor of modern systems (50).

Based on the determined intracellular concentration of 2OG in *M. maripaludis*, ranging from 0.08 mM under non-limiting conditions to 0.8 mM 2OG under nitrogen starvation (15–17), we propose that the post-translational regulatory response is sequential in *Methanococcales*. They would first respond to nitrogen limitation by deactivating the P_II_-protein GlnK (e.g., with an affinity for 2OG estimated in the µM range (27, 36)), enhancing NH_3_ import. Then, NH_3_ fixation would be stimulated via 2OG-dependent allosteric activation of glutamine synthetase (i.e., K_0.5_ to be 0.17 mM (46)). Finally, since the K_0.5_ of NifI_1,2_ is in the mM range for 2OG, nitrogenase would be used only if cellular nitrogen levels are critically low and constitutes the “last resort” for nitrogen acquisition (16, 51).

To conclude, by unveiling the N_2_-fixation inhibition mechanism, we demonstrate how NifI_1_ and NifI_2_ form a higher oligomerisation state to specifically bind to the nitrogenase, while conserving the universal principle of a functional switch triggered by the cellular energy charge and nitrogen availability (Fig. 4). The T-loop conformational switch imposed by ligand binding is provoking steric clashes and disrupts contacts with the target, as previously observed in other P_II_-family proteins (e.g., GlnK-AmtB inhibition mechanism (19, 26)), illustrating a conserved mechanistic with a unique target-binding mode due to an original oligomerisation. The observed sequestration phenomenon by NifI_1,2_ blocks unnecessary cellular energy loss and preserves the nitrogenase in a standby mode rather than degrading it, thereby conserving the substantial energy invested in the biosynthesis of the metallocofactors (52). Because of this energy-saving main function, it is reasonable to argue that the predominant presence of NifI_1,2_ in anaerobes reflects the energy limitation of these microbes compared to aerobes. By deriving additional energy gain through oxygen respiration, aerobes might have progressively lost NifI_1,2_ to rather focus on accessory proteins as strategies to protect the N_2_-fixation apparatus from oxidative damage by direct protection (e.g., via the Shethna protein II (Supplementary Fig. 2a)), transcriptional regulation (e.g., via the NifLA-GlnK system that regulates the expression of nitrogen fixation genes in response to intracellular oxygen, cellular energy, and the carbon–nitrogen ratio (53, 54)), and deactivation of O_2_-highly sensitive NifH (e.g., by DraT that inactivates NifH through ADP-ribosylation when ammonium is abundant, and DraG, which removes the ribosylation under nitrogen-limiting conditions (55, 56)). The latter two strategies have been reported to be under the control of P_II_-proteins (57, 58). Combining the knowledge acquired in this work with bioengineering rationale developed for the P_II_-family (36) could lead to future studies targeting the ligand-binding site of NifI_1,2_ to alter 2OG sensing or redesign it for analogue recognition. With these molecular tools, it would become feasible to switch on and off, and tweak N₂-fixation rates in a methanogenic or acetogenic chassis engineered through synthetic biology, enabling H₂-driven green carbon cycling (e.g., methane, ethanol, or 2,3-butanediol production from CO₂) alongside NH₃ bioproduction.

## Methods

### Microbial growth conditions

All proteins were natively isolated from *Methanocaldococcus infernus* strain DSM 11812 (Leibniz Institute DSMZ - German Collection of Microorganisms and Cell Cultures, Braunschweig, Germany). The anaerobic medium was prepared as in Maslać *et al.* 2025 (31), and cells were cultivated in batches in 1-L Duran bottles. Diazotrophic growth conditions, harvesting, and cell storage were handled as previously described (31). *M. infernus* was routinely cultivated anaerobically in 30-60 mL medium in 1-L pressure-resistant Duran bottles with 0.5-2 mM final concentration of Na_2_S as a reductant and sulfur source. The anaerobic medium was transferred to a Duran bottle in a glovebox under a N_2_/CO_2_ (90:10%) atmosphere and sealed with a rubber stopper. Once transferred out, the headspace of the bottle was replaced by first vacuuming to −1 bar, then filling to 0.5 bar H_2_/CO_2_ (80:20%) and complementing to 1 bar with pure N_2_. The culture was started by adding the inoculum in a 1:20 ratio and was incubated standing without agitation at 75 °C in the dark. After roughly 24 h, the gas phase was refreshed using the same procedure of vacuuming followed by H_2_/CO_2_ and N_2_ addition. When the culture reached a late exponential phase on the second day (i.e., when OD_600nm_ reached 1.0–1.5), cells were transferred to an anaerobic glove box containing a gas mixture of N_2_/CO_2_ (90:10%). Cells were harvested anaerobically at room temperature by centrifugation at 6,000 x *g* for 30 min at 20 °C. The wet cell pellet was placed in a sealed serum flask and anaerobically transferred out of the glove box, where 0.8 bar of N_2_ was added on top. The flask was finally stored at −80 °C.

### Protein purification

The native purification of NifDKI_1,2_, NifDK, NifH, and NifI_1,2_ was successfully performed several times using similar protocols. The purification was monitored by denaturing gel electrophoresis and by following absorbance at 280, 415, and 550 nm.

27.21 g of *M. infernus* cell pellets obtained from a total of 5.5 L of culture were thawed at room temperature inside a glovebox under a N_2_/CO_2_ (90:10%) atmosphere. Six volumes of lysis buffer (50 mM Tris/HCl, pH 8.0, 2 mM dithiothreitol (DTT)) were added, and the cell resuspension was sonicated (Bandelin SONOPULS, Berlin, Germany, KE76 probe at 75% intensity). The lysate was centrifuged at 45,000 x *g* for 60 minutes at 25 ⁰C and transferred to an anaerobic Coy tent filled with an N_2_/H_2_ (97:3%) atmosphere. The whole purification procedure was performed under yellow light at 20 °C. The supernatant was filtered through a 0.2 μm filter (Sartorius, Göttingen, Germany) and loaded on a 10 mL HiTrap Q-Sepharose High-Performance column (Cytiva, Freiburg, Germany) previously equilibrated with lysis buffer. The sample was eluted over a 250 to 700 mM NaCl linear gradient for 5 column volumes (CVs) at a 1.5 mL/min flow rate. NifDKI_1,2_ was eluted between 360 and 514 mM NaCl. NifH eluted between 250 and 360 mM NaCl and was kept aside for further purification following the procedure described previously (31).

One volume of hydrophobic exchange buffer (25 mM Tris/HCl pH 8.0, 2 M (NH_4_)_2_SO_4_ and 2 mM DTT) was added to NifDKI_1,2_ pool. The sample was filtered on a 0.2 µm filter (Sartorius, Göttingen, Germany) and immediately loaded twice at 1.5 mL/min on a 1.7 mL Source 15PHE 4.6/100 column (Cytiva, Freiburg, Germany) pre-equilibrated with the hydrophobic exchange buffer. NifDKI_1,2_ was eluted with 1-0.3 M (NH_4_)_2_SO_4_ linear gradient for 23.5 CV at a 1 mL/min flow rate. The protein eluted between 0.79 and 0.51 M (NH_4_)_2_SO_4_.

Part of the NifDKI_1,2_ pool (usually half or a third) was diluted with three volumes of lysis buffer, filtered through a 0.2 µm filter, and loaded on a 5 ml HiTrap Q-Sepharose High-Performance column (Cytiva, Freiburg, Germany) previously equilibrated with lysis buffer. The elution was performed with a linear gradient from lysis buffer to 500 mM sodium malate pH 5.4, and 2 mM DTT (buffered with KOH pellets, Merck, Darmstadt, Germany). Under these conditions, NifDKI_1,2_ eluted between 219.85 and 272.2 mM malate.

The remaining NifDKI_1,2_ pool (usually half or two-thirds) was used to purify NifDK and NifI_1,2_. First, the complex NifDK-NifI_1,2_ was dissociated by incubating the sample in lysis buffer supplemented with fresh 2 mM ATP, 2 mM MgCl_2_, and 10 mM 2OG final (all components were freshly added anaerobically) for a few hours to an overnight. Then, the sample was passed through a 0.2 µm filter and injected on a 1 mL HiTrap Q-Sepharose High-Performance column (Cytiva, Freiburg, Germany) previously equilibrated with lysis buffer containing 2 mM ATP, 2 mM MgCl_2_, and 10 mM 2OG. Depending on the protein quantity, the procedure can be done all at once or twice by splitting the injected sample. The elution was performed at 0.5 mL/min with a NaCl gradient from 100 to 600 mM in 60 CV. NifDK eluted at 215-310 mM NaCl and NifI_1,2_ at 385-455 mM NaCl.

To obtain the stable NifDK-NifH complex, a molar ratio of 2 isolated NifH for 1 isolated NifDK was mixed and incubated for 20 minutes at room temperature in the following buffer: 25 mM Tris/HCl pH 8.0, 20 mM NaF, 1 mM AlCl_3_, 1 mM ADP, and 2 mM MgCl_2_. This sample was filtered and loaded on a 5 mL HiTrap Q-Sepharose High-Performance column (Cytiva, Freiburg, Germany) equilibrated with the same buffer.

Elution at 1.5 mL/min was performed with a gradient ranging from 100 to 600 mM NaCl over 10 CV. Under this condition, the NifDKH complex eluted between 109.10 and 377.20 mM NaCl.

Finally, NifI_1,2_ was further purified by hydrophobic interaction chromatography. Two volumes of the hydrophobic exchange buffer were added to the NifI_1,2_ pool, and the sample was filtered through a 0.2 µm filter. The filtrate was immediately loaded at 1.0 mL/min on a Source 15PHE 4.6/100 column (Cytiva, Freiburg, Germany) pre-equilibrated with the hydrophobic exchange buffer. NifI_1,2_ was eluted with a 1.4-0 M (NH_4_)_2_SO_4_ linear gradient for 28 CV at 0.8 mL/min. Under these conditions, the protein eluted between 0.5 and 0.33 M (NH_4_)_2_SO_4_.

As a comment, the fact that the functional unit inhibiting the nitrogenase is composed of both NifI_1_ and NifI_2_ explains why overexpression of tagged NifI_1_ or NifI_2_ alone does not promote NifDK capture (16). We previously showed that the transcript level of NifI_2_ is similar to that of NifI_1_, suggesting that the cell controls the assembly of a functional complex, rather than the inactive homotrimeric NifI_1_ or homotrimeric NifI_2_, at the translational level. Based on the soluble fraction profile and the fact that an unbound population of NifI_1,2_ is detected while native NifDK is fully saturated with NifI_1,2_ (31), it is assumed that the cytoplasmic level of NifI_1,2_ is always superior to that of NifDK. With this, NifI_1,2_ would constitutively be present to immediately shut down N_2_-fixation by blocking NifH binding on the nitrogenase, preventing unnecessary ATP-consumption if the cellular nitrogen is sufficient (Fig. 4).

### Crystallisation and structure determination

All samples were concentrated by centrifugation using VivaSpin Amicons (Sartorius, Göttingen, Germany). Concentrators with 3 kDa, 10 kDa, and 30 kDa cut-offs were used for NifI_1,2_, NifH, and NifDKI_1,2_/NifDKH, respectively. Samples were further centrifuged at 13,000 x *g* for 3 min to remove macroaggregates and dust. All protein mixtures were crystallised in an anaerobic Coy tent filled with an N_2_/H_2_ (97:3%) atmosphere at 20 °C. Crystallisation was performed by the sitting-drop method in 96-Well MRC 2-Drop polystyrene crystallisation plates (SWISSCI), with 90 μL of crystallisation solution in the reservoir in all cases.

NifDKI_1,2_ was crystallised at a final concentration of 26.7 mg/mL by spotting 0.6 μL of crystallisation solution with 0.6 μL of protein sample. The crystallisation solution contained 2 M (NH_4_)_2_SO_4_, 0.1 M Sodium HEPES pH 7.5. The crystal shape was a brown polygon that took several months to appear. Before freezing in liquid nitrogen, the crystals were soaked in 2 M (NH_4_)_2_SO_4_, 0.1 M Bis-Tris pH 5.5, supplemented with 30% (v/v) glycerol for a few seconds.

NifI_1,2_ holo was crystallised aerobically at a final concentration of 15.2 mg/mL supplemented with 2 mM ATP, 2 mM MgCl_2_ and 10 mM 2OG. Crystals were prepared by spotting 0.5 μL of crystallisation solution with 0.5 μL of protein sample, and were obtained in the solution containing 30% (v/v) polyethylene glycol 400, 100 mM Tris pH 8.5 and 200 mM magnesium chloride. Small stacked transparent plates appeared after a few months.

NifDKH was crystallised at a final concentration of 17.2 mg/mL with 20 mM NaF, 1 mM AlCl_3_, 1 mM ADP, and 2 mM MgCl_2_ by spotting 0.5 μL of crystallisation solution with 0.5 μL of protein sample. Crystals were obtained in the solution containing 10% (w/v) polyethylene glycol 8,000, 100 mM Tris pH 7.0, and 200 mM MgCl_2_. Before freezing in liquid nitrogen, the crystals were soaked in the crystallisation solution supplemented with 30% (v/v) glycerol for a few seconds. The crystals had a brown rod morphology and appeared within a few weeks.

### Data collection and structural analysis

The diffraction experiments used for the deposited model were performed at 100 K on beamlines BM07-FIP2 at the European Synchrotron Radiation Facility (ESRF), X06-SA PXI at the Swiss Light Source synchrotron (SLS), and PROXIMA-2 at the “Source optimisée de lumière d’énergie intermédiaire du LURE” (SOLEIL synchrotron, see Supplementary Table 1).

The data were processed and scaled with autoPROC (59). The X-ray diffraction showed a slight anisotropy for NifDKI_1,2_ and NifI_1,2_ holo and a pronounced one for NifDKH. Hence, we used the STARANISO correction integrated with the autoPROC pipeline (60) (STARANISO. Cambridge, United Kingdom: Global Phasing Ltd.).

The structure of NifI_1,2_ was solved using a computationally generated AlphaFold 2 model as a template for molecular replacement (61). The model was then used in the NifDKI_1,2_ supercomplex for molecular replacement together with the NifDK atomic-resolution structure previously obtained in Maslać *et al*. (2025)(31). The same NifDK atomic-resolution model (31) was used for molecular replacement in the NifDKH structure, together with NifH from *M. infernus* obtained with MgADP (37). Because of the extremely high translational non-crystallography symmetry (i.e., 33% fraction) in NifDKH and the wobbly state of the second bound NifH dimer, only the most stable NifH dimer was manually refined, and its model was docked on the wobbly position without allowing xyz-displacement over real-space refinement. In all cases, molecular replacement was done with PHASER from the Phenix package (62, 63).

Models were manually built via Coot (64) and refined with phenix.refine (62). The refinement steps were performed by applying a translational-libration-screw without non-crystallographic symmetry and without modelling hydrogens. Models were validated by the MolProbity server (65). Data collection and refinement statistics for the deposited models are listed in Supplementary Table 1. All figures with structures were generated and rendered with PyMOL (Version 2.2.0, Schrödinger, LLC, New York, NY, United States).

Sequence alignment was performed with MUSCLE (66) from the EBI and Supplementary Figs. 3 and 4 were generated with the Espript server (67).

Computational models in Fig. 4 were all generated by the AlphaFold 3 server (68). Since the modelled T-loops conflicted with the dimeric arrangement, T-loops from NifI_1_ and NifI_2_ sequences were removed from all queries.

## Data Availability

NifDKI_1,2_, NifI_1,2_ holo and NifDKH structures were validated and deposited in the Protein Data Bank (PDB) under the following accession numbers: 9TJ1, Crystal structure of the nitrogenase from *Methanocaldococcus infernus* inhibited by the P_II_ proteins NifI_1,2_. 9TIZ, Crystal structure of the P_II_ proteins from *Methanocaldococcus infernus* NifI_1,2_ bound to MgATP and 2-oxoglutarate. 9TJ2, Crystal structure of the nitrogenase from *Methanocaldococcus infernus* bound to NifH complexed with MgADP-AlF_4_ ^-^.

## Author contributions

Organism cultivation, protein purification, and crystallisation were done by N.M., P.B. and T.W. X-ray diffraction experiments were performed by all authors. N.M., M-C.M. and T.W. performed the data processing, model building, and structure refinement. T.W. validated and analysed the structures. T.W. wrote the manuscript with contributions and final approval of all co-authors.

## Competing interest statement

Authors declare no competing interests.

## Acknowledgements

We thank the Max Planck Institute for Marine Microbiology and the Max Planck Society for their continuous support. We also thank Christina Probian and Ramona Appel for their continuous technical support in the Microbial Metabolism laboratory. We acknowledge the French Biology/Health Panel Review Committee for providing synchrotron radiation beamtime at the ESRF (Grenoble, France) on beamline BM07-FIP2. BM07-FIP2 is supported by the French ANR PIA3(France 2030) EquipEx+ project MAGNIFIX under grant agreement ANR-21-ESRE-0011. We thank the SLS for beamtime allocation and the staff of the beamline X06SA. We also thank the beamtime allocation for the beamline at SOLEIL and the local support on PROXIMA II at SOLEIL and BM07-FIP2 at the ESRF.

---

**Supplementary Table 1.** Data collection and refinement statistics.

|  | NifDKI<sub>1,2</sub> | NifI<sub>1,2</sub><br>holo | NifDKH<br>with MgADP-AlF<sub>4</sub><sup>-</sup> |
| --- | --- | --- | --- |
| **Data collection** |  |  |  |
| Synchrotron, beamline | ESRF, BM07-FIP2 | SOLEIL, PROXIMA II | SLS, X06SA (PXI) |
| Wavelength (Å) | 0.97951 | 0.97625 | 1.00000 |
| Space group | *P*6<sub>3</sub> | *P*3<sub>1</sub>2<sub>1</sub> | *P*2<sub>1</sub>2<sub>1</sub>2<sub>1</sub> |
| Resolution (Å) | 146.73 – 2.27<br>(2.49 – 2.27) | 56.14 – 2.70<br>(2.77 – 2.70) | 48.93 – 2.59<br>(2.82 – 2.59) |
| Cell dimensions |  |  |  |
| a, b, c (Å) | 293.46, 293.46, 108.65 | 68.96, 68.96, 164.81 | 100.31, 154.75, 222.41 |
| α, β, γ (°) | 90, 90, 120 | 90, 90, 120 | 90, 90, 90 |
| R<sub>merge</sub> (%)<sup>a</sup> | 11.2 (175.2) | 13.2 (166.3) | 38.1 (135.5) |
| R<sub>pim</sub> (%)<sup>a</sup> | 3.4 (53.1) | 4.1 (49.9) | 13.2 (47.7) |
| CC<sub>1/2</sub><sup>a</sup> | 0.992 (0.558) | 0.993 (0.594) | 0.995 (1.000) |
| I/σ<sub>I</sub><sup>a</sup> | 12.6 (1.5) | 9.9 (1.6) | 5.0 (1.6) |
| Spherical completeness (%)<sup>a</sup> | 77.7 (16.3) | 96.8 (61.2) | 43.4 (9.6) |
| Ellipsoidal completeness (%)<sup>a</sup> | 96.2 (66.8) | 96.8 (61.2) | 89.2 (63.5) |
| Redundancy<sup>a</sup> | 11.7 (11.8) | 11.1 (11.3) | 9.1 (8.8) |
| Nr. unique reflections<sup>a</sup> | 190,910<br>(9,545) | 12,691<br>(635) | 47,071<br>(2,354) |
| **Refinement** |  |  |  |
| Resolution (Å) | 27.58 – 2.27 | 56.14 – 2.70 | 48.93 – 2.59 |
| Number of reflections | 190,693 | 12,687 | 47,002 |
| R<sub>work</sub>/R<sub>free</sub><sup>b</sup> (%) | 17.00/19.17 | 23.14/26.04 | 21.82/26.01 |
| Number of atoms |  |  |  |
| Protein | 20,097 | 2,454 | 23,367 |
| Ligands/ions | 1,058 | 220 | 469 |
| Solvent | 543 | 15 | 189 |
| Mean B-value (Å<sup>2</sup>) | 77.43 | 80.16 | 34.14 |
| Molprobity clash<br>score, all atoms | 2.29 | 0.56 | 4.16 |
| Ramachandran plot |  |  |  |
| Favoured regions (%) | 96.82 | 97.08 | 96.62 |
| Outlier regions (%) | 0 | 0 | 0 |
| rmsd<sup>d</sup> bond lengths (Å) | 0.006 | 0.012 | 0.010 |
| rmsd<sup>d</sup> bond angles (°) | 0.797 | 1.566 | 1.254 |
| **PDB ID code** | 9TJ1 | 9TIZ | 9TJ2 |

<sup>a</sup> Values relative to the highest resolution shell are within parentheses. <sup>b</sup> R<sub>free</sub> was calculated as the R<sub>work</sub> for 5% of the reflections that were not included in the refinement.

**Supplementary Fig. 1:**
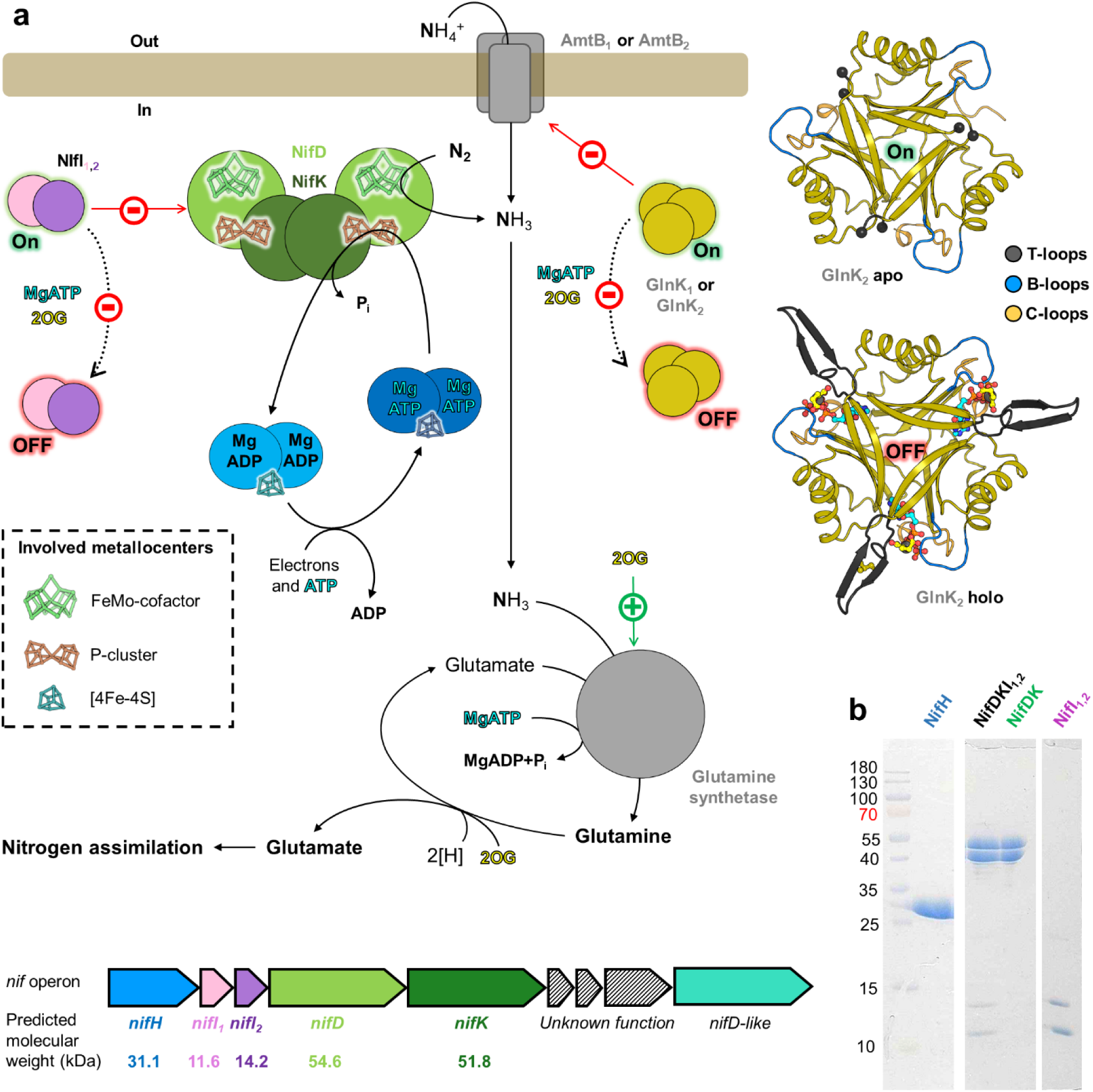
Actors of the nitrogen assimilation in *Methanococcales*. **a**, Overall scheme of the nitrogen assimilation in *Methanococcales* gathered from previous work (16, 27, 31, 46, 69) and *nif* operon organisation in *M. infernus* (bottom). The structures of GlnK_2_ from *Methanothermococcus thermolithotrophicus* apo (PDB 7P4Y) and holo (PDB 7P50) are presented on the top right. The homotrimers are shown in cartoons with the ligand-binding loops coloured differently. Mg^2+^ (grey), ATP (cyan carbon), and 2-oxoglutarate (2OG, yellow carbon) are displayed in balls and sticks. Glutamate production from glutamine and 2OG is catalysed by the glutamate synthase (GOGAT). **b.** Denaturing gels of NifH, NifDKI_1,2_, NifDK, and NifI_1,2_ native purification.

**Supplementary Fig. 2:**
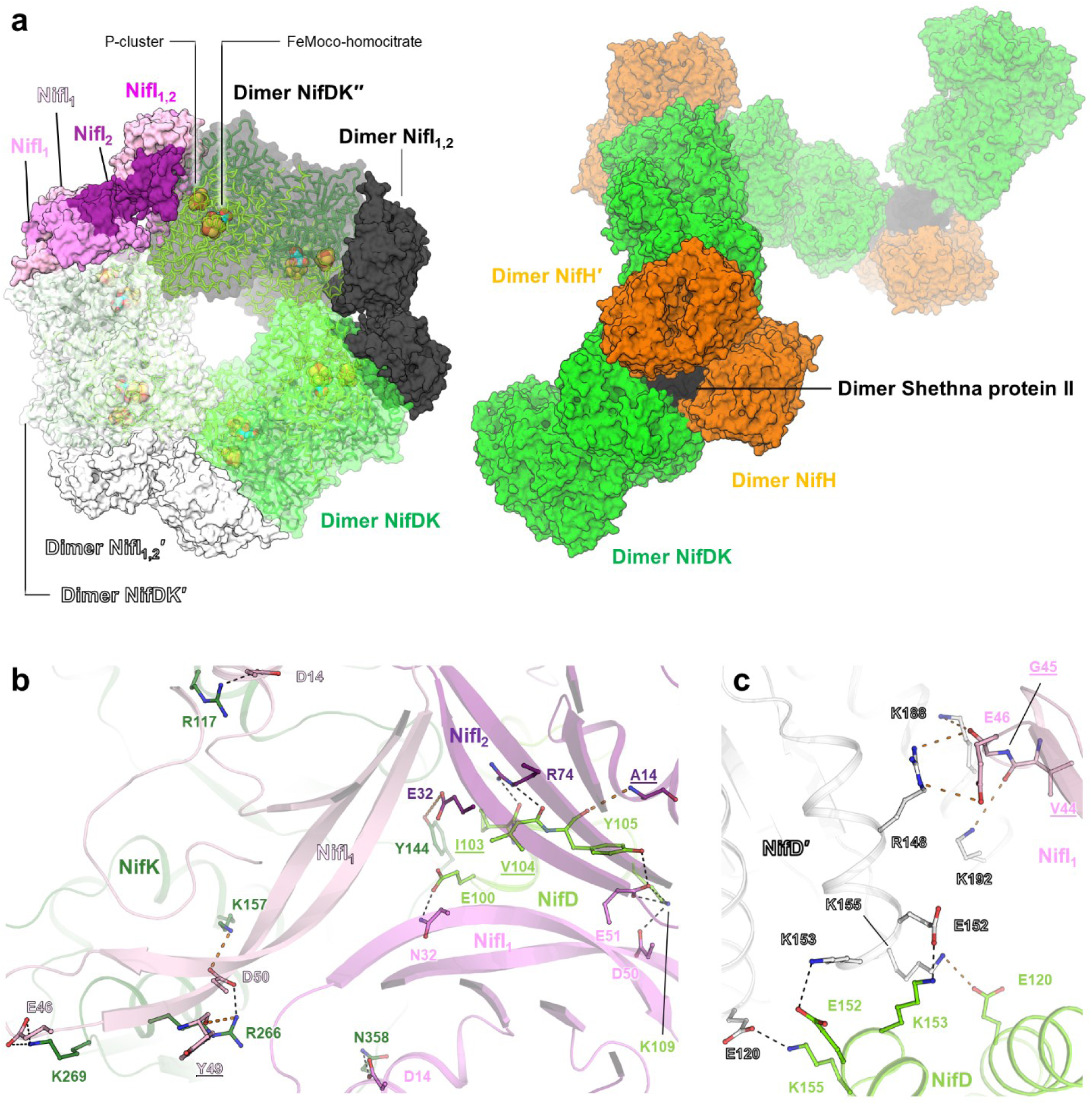
Comparisons of NifI_1,2_-inhibited NifDK with Shethna-protected NifDK and details of NifI_1,2_ binding site. **a**, Overall NifDKI_1,2_ assembly highlighting the metallocofactor position. Compared to Fig. 1a, NifI_1,2_ is displayed as surfaces, and NifDK as transparent surfaces with metallocofactors highlighted as balls. The right panel presents the structure of the NifDK and NifH complex from *Azotobacter vinelandii* protected by the Shethna protein II (PDB 8RHP). Here, the complex is represented as a surface and organised into fibrils. **b**, NifI_1,2_-NifDK, and **c**, NifD-NifD binding contacts in the inhibited supercomplex. Proteins are shown as cartoons with residues interacting via hydrogen bonds (displayed by dashed lines) highlighted as sticks. Underlined residues interact solely via the main chain.

**Supplementary Fig. 3:**
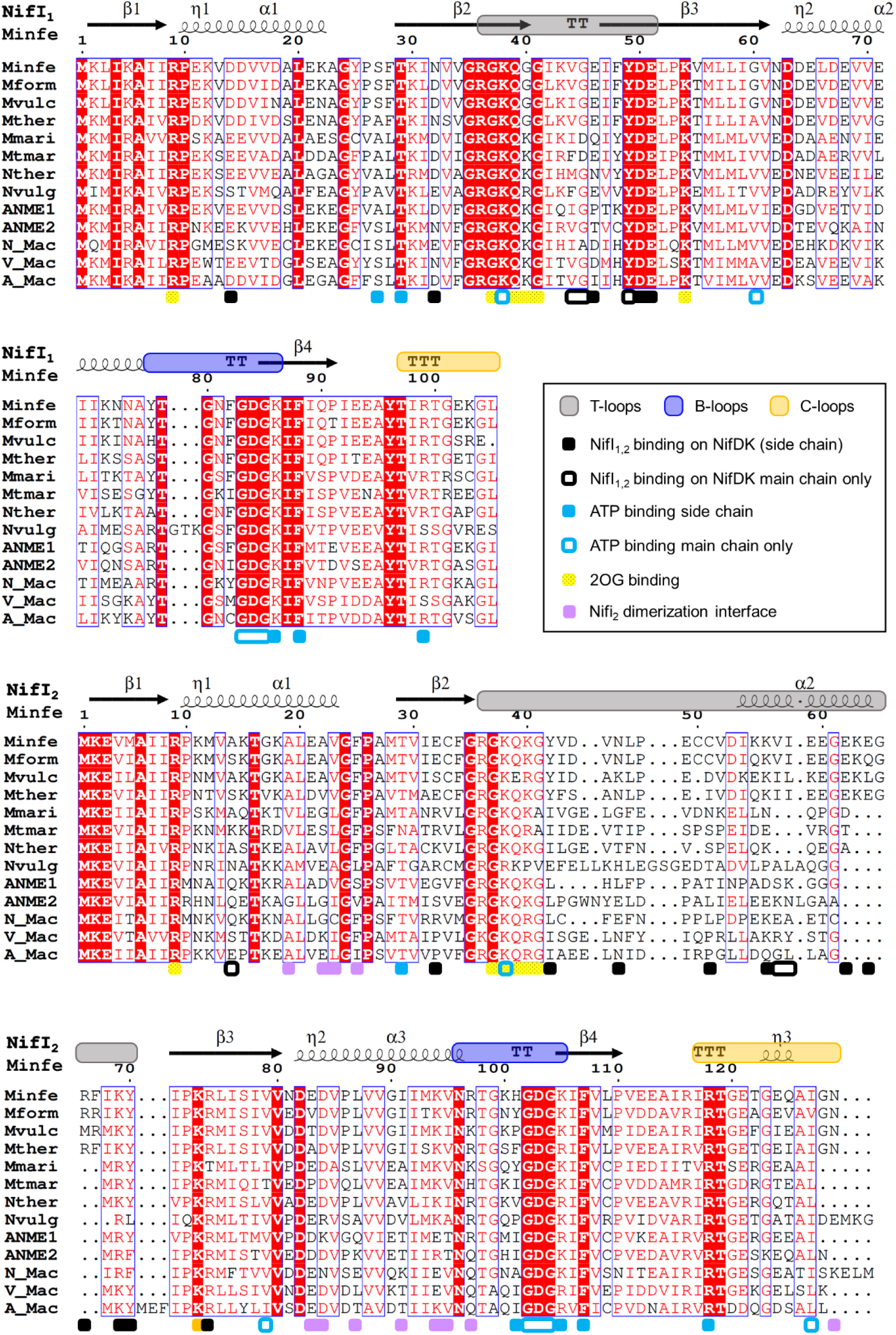
Conservation in NifI_1_ and NifI_2_. Full organism names are provided in the Suppl. Fig. 4. Sequence alignment superposed on the secondary structure was performed with Espript (67).

**Supplementary Fig. 4:**
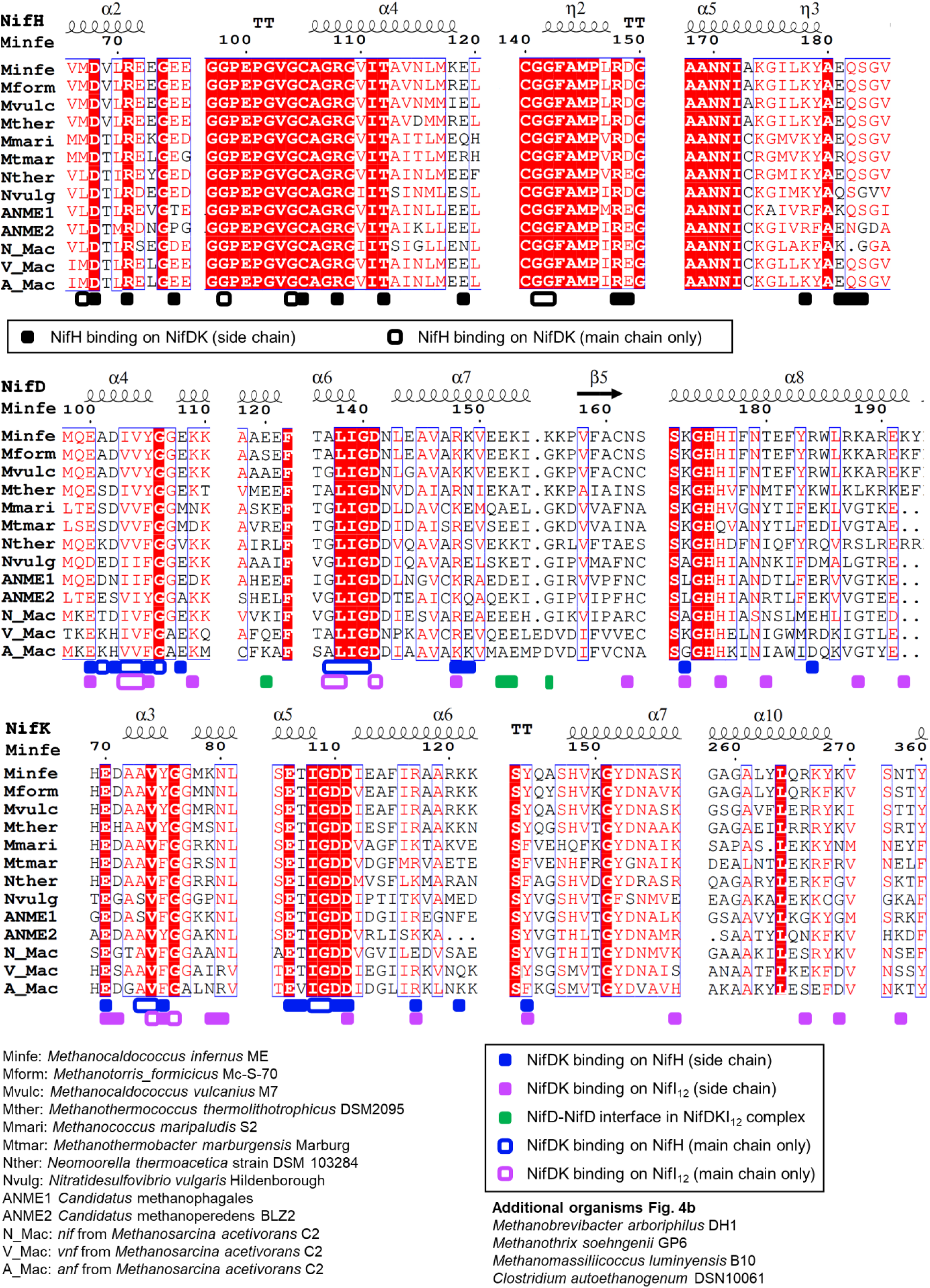
Conservation in NifH, NifD, and NifK. Sequence alignment superimposed on the secondary structure was performed with Espript (67).

**Supplementary Fig. 5:**
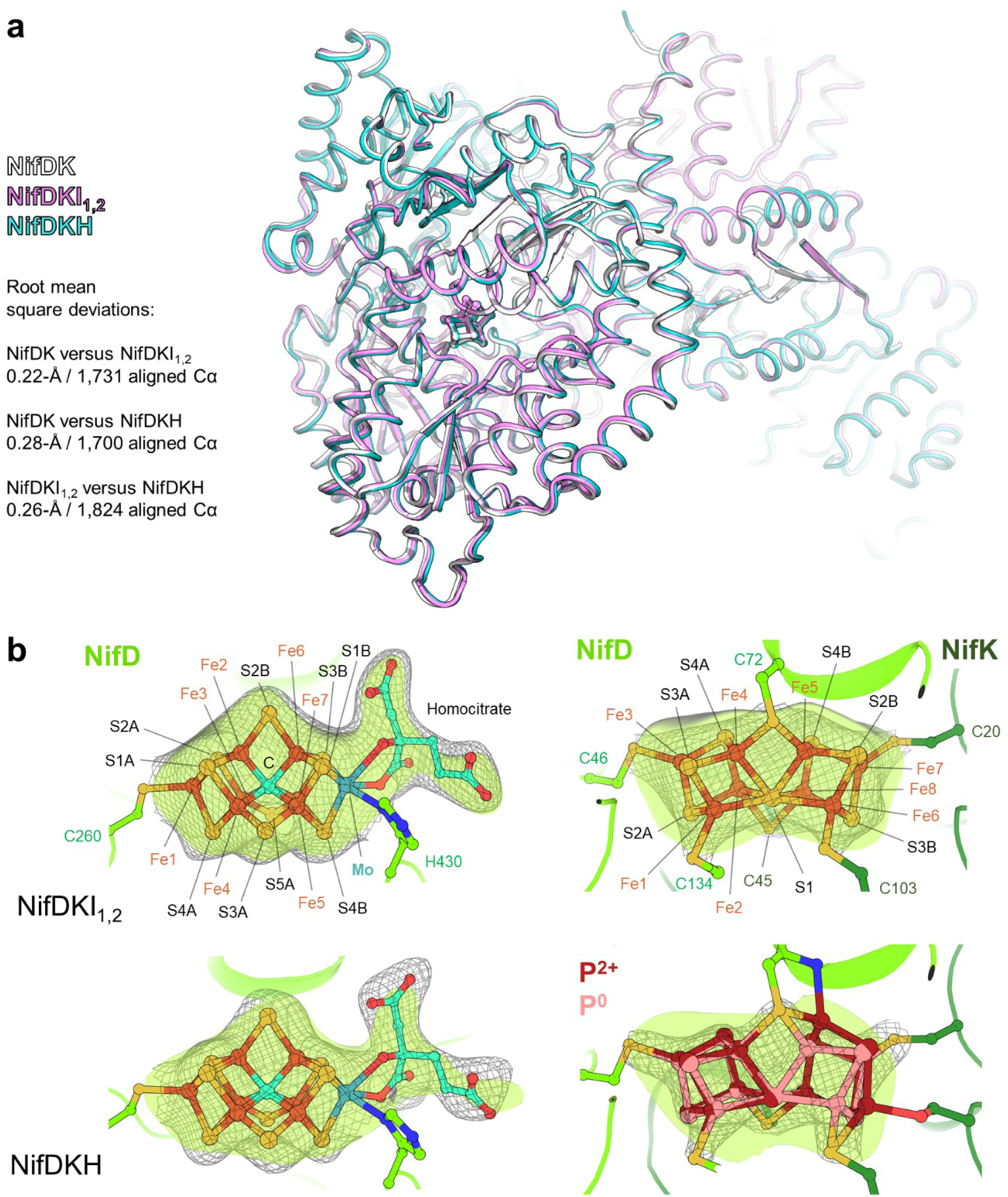
Overall comparison between the different trapped complexes and metallocofactor sites of NifDK. **a**, overall superposition of *M. infernus* NifDK (PDB 9SQ3 (31)), NifDKI_1,2_ and NifDKH. Root mean square deviations are indicated. **b**, Close up of the FeMo-cofactor (left) and P-cluster (right) binding sites with ligands and protein-coordinating residues shown as balls and sticks. The 2*F*_o_−*F*_c_ maps are displayed as grey mesh and contoured at 1-σ for the FeMo-cofactor/homocitrate, and 4-σ for the P-cluster. The positive difference maps resulting from the refined model omitting the metallocofactor are displayed as a transparent green surface and contoured at 3.5-σ in all panels.

**Supplementary Fig. 6:**
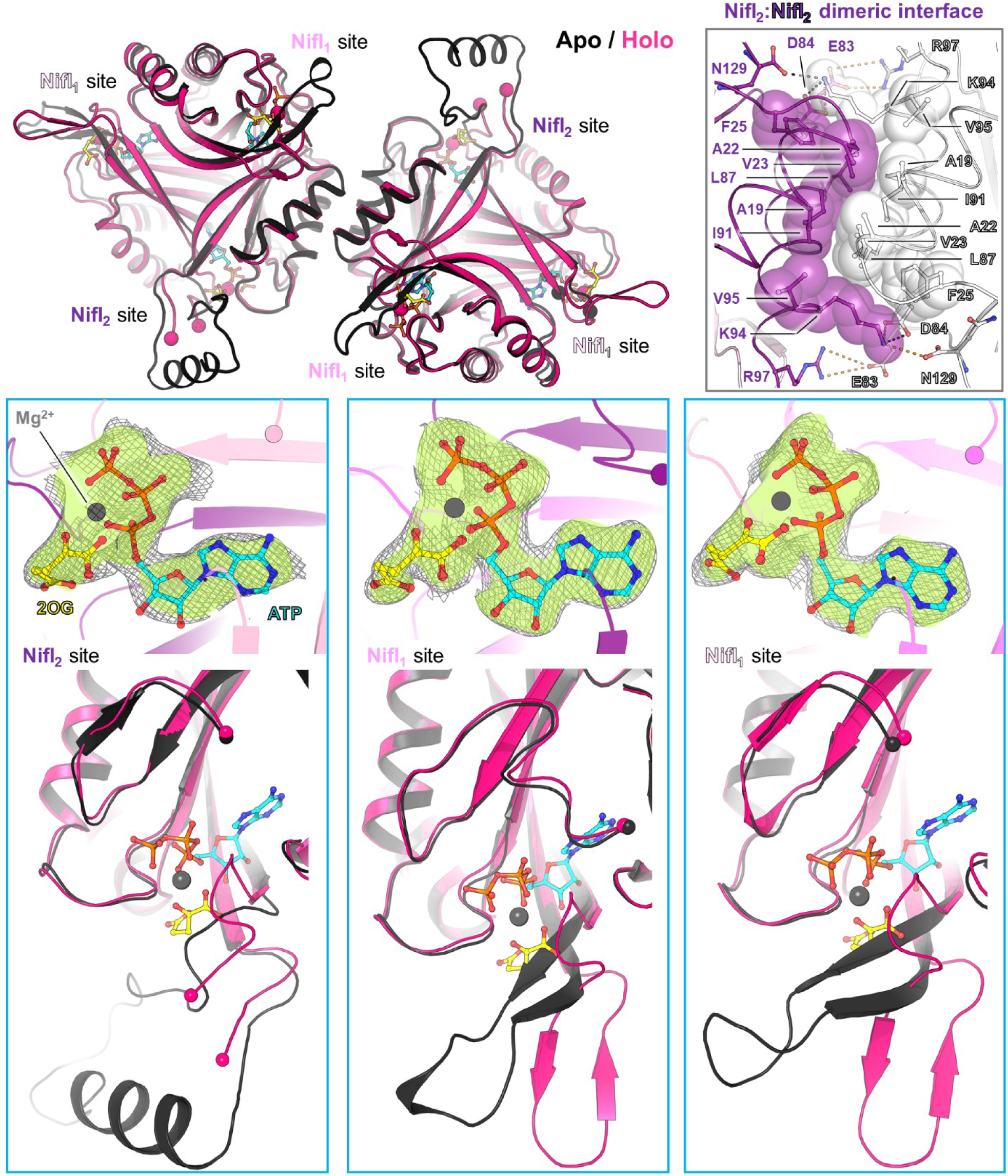
Superposition of apo and holo NifI_1,2_. The top-left panel presents an overall superposition of the NifI_1,2_ models in cartoons. The top-right panel shows a close-up of the dimeric interface, with residues involved in salt bridges (dashed lines) and Van der Waals interactions shown as sticks and balls, respectively. Bottom panels show the electron density of the ligands and a close-up of the superposition on the main chain binding the ligands. The 2*F*_o_−*F*_c_ maps are displayed in grey mesh and contoured at 1-σ. The positive difference maps resulting from the refined model omitting the ligands are displayed as a transparent green surface and contoured at 2.7-σ in all panels. All ligands are represented as balls and sticks.

**Supplementary Fig. 7:**
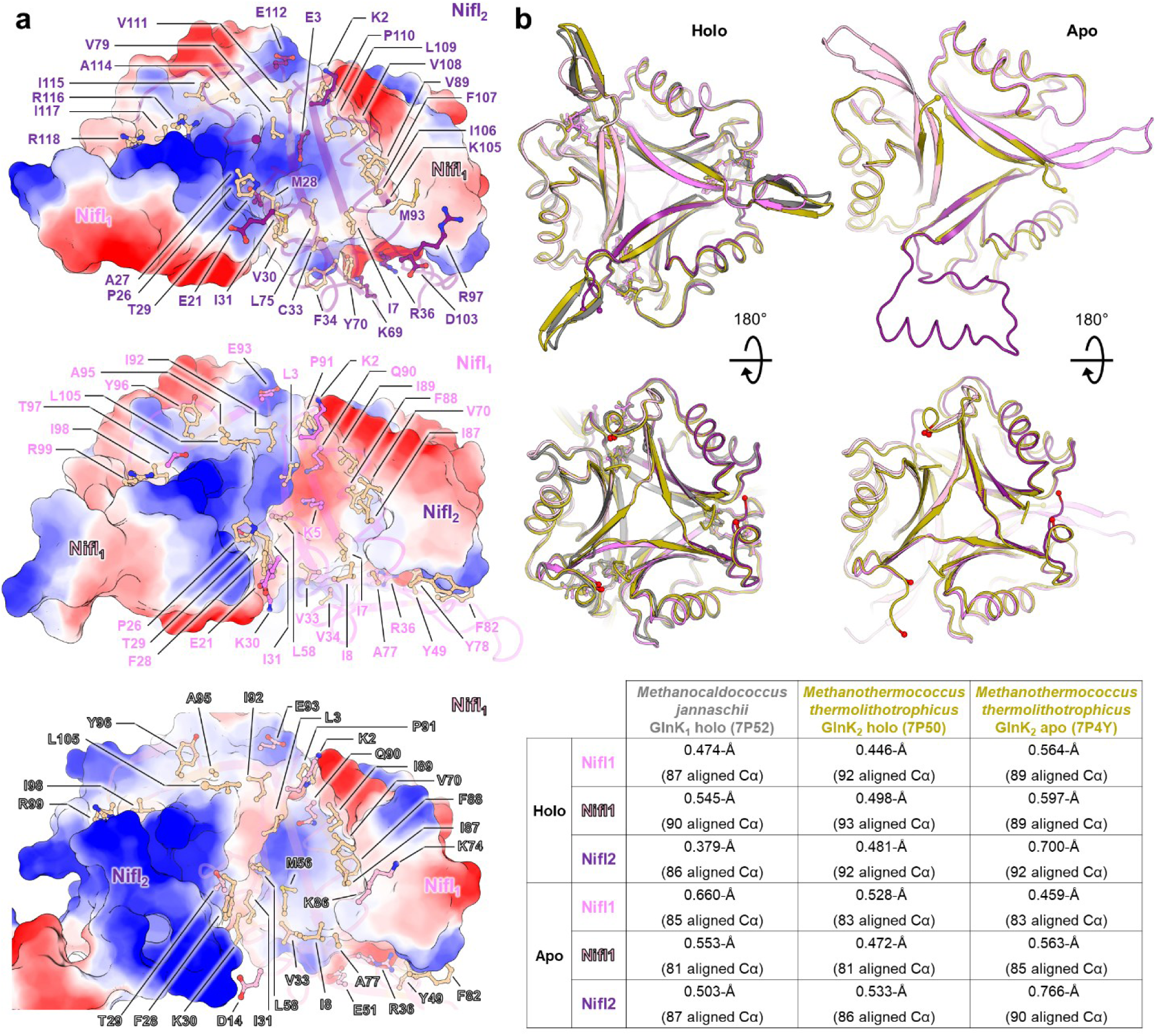
NifI_1,2_ intramolecular organisation and comparison with other P_II_–family proteins. **a**, Electrostatic surface complementation between one selected NifI subunit (displayed as transparent cartoons) and the two others (shown in surface), ranging from negatively (red) to positively (blue) charged. Residues involved in hydrogen bonds are shown as sticks with carbons coloured by the subunit code (i.e., violet, pink, light pink), whereas residues involved in Van der Waals interactions are colored wheat. **b**, Superposition of GlnKs from *M. jannaschii* and *M. thermolithotrophicus* with NifI_1,2_. The superposition has been done on NifI_2_. The colour codes and root-mean-square deviations are reported in the table.

**Supplementary Fig. 8:**
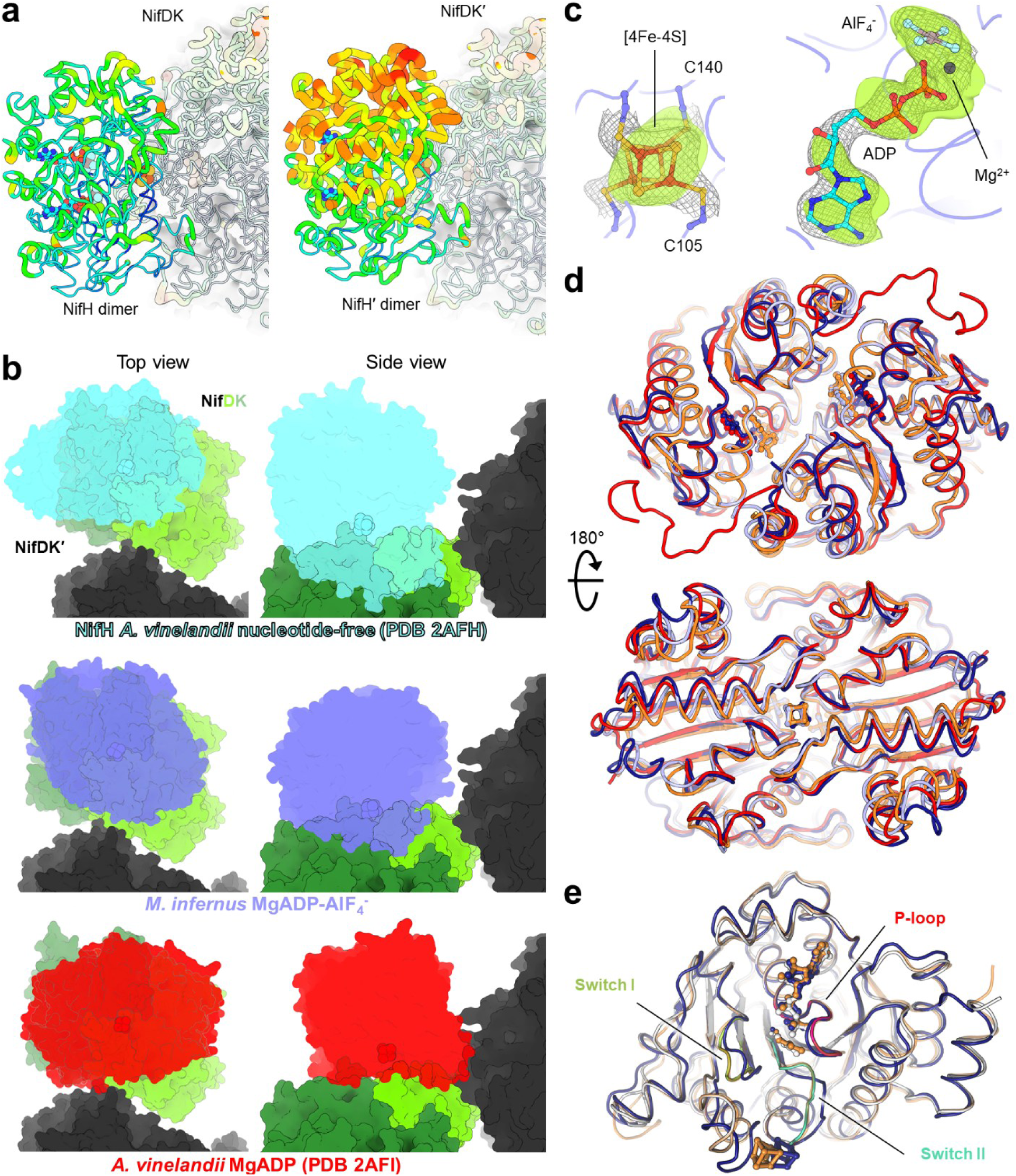
NifH state and binding position. **a**, B-factor comparison (low values in blue and thin to high values red and thick) between both NifH dimers occupying the asymmetric unit. NifDK is covered by a white transparent surface. **b**, Comparison of NifH position (transparent surface) on the trimeric NifDK arrangement imposed by NifI_1,2_, in which NifDK is colored in green, and NifDK′ in black. **c**, 2*F*_o_−*F*_c_ (contoured at 1-σ) and omit maps (contoured at 2.7-σ) are displayed in grey mesh and transparent green surface, respectively. **d**, Overall superposition of *M. infernus* MgADP-NifH (navy, PDB 8Q5W), and with MgADP-AlF_4_^-^ (light blue), *A. vinelandii* MgADP-NifH (red, PDB 1FP6) and with MgADP-AlF_4_^-^ (orange, PDB 1M34). **e**, Single chain superposition of *M. infernus* MgADP-NifH (navy) and with MgADP-AlF_4_^-^(white with colored loops), and *A. vinelandii* MgADP-AlF_4_^-^ (orange). Ligands are in balls and sticks.

**Supplementary Fig. 9:**
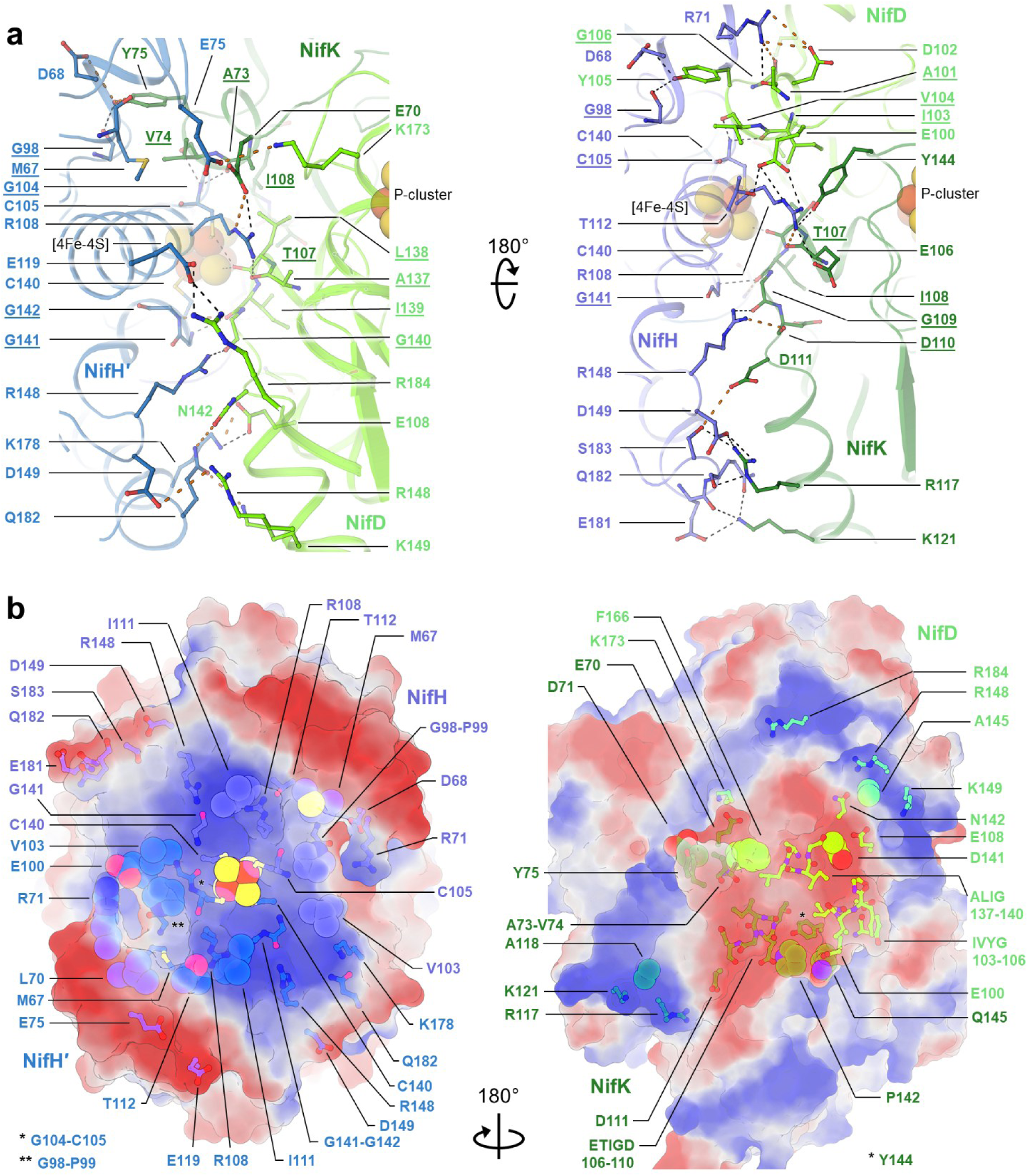
NifH-NifDK interaction. **a**, Close-up of the interaction of NifH with NifDK. Proteins are shown in cartoons with residues interacting via hydrogen bonds (displayed by dashed lines) highlighted in sticks. Dashed lines in orange are long distances comprised between 3.5-4.0 Å. **b**, Electrostatic surface complementation between dimeric NifH (left) and NifDK (right), ranging from negatively (red) to positively (blue) charged. Residues involved in hydrogen bonds and Van der Waals interactions are shown in sticks and balls, respectively.

## Notes

### Competing Interest Statement

The authors have declared no competing interest.

